# Deep generative model embedding of single-cell RNA-Seq profiles on hyperspheres and hyperbolic spaces

**DOI:** 10.1101/853457

**Authors:** Jiarui Ding, Aviv Regev

## Abstract

Single-cell RNA-Seq (scRNA-seq) has become an invaluable tool for studying biological systems in health and diseases. While dimensionality reduction is a crucial step in interpreting the relation between cells based on scRNA-seq, current methods often are hampered by “crowding” of cells in the center of the latent space, biased by batch effects, or inadequately capture developmental relationships. Here, we introduced scPhere, a scalable deep generative model to embed cells into low-dimensional hyperspherical or hyperbolic spaces, as a more accurate representation of the data. ScPhere resolves cell crowding, corrects multiple, complex batch factors, facilitates interactive visualization of large datasets, and gracefully uncovers pseudotemporal trajectories. We demonstrate scPhere on six large datasets in complex tissue from human patients or animal development, demonstrating how it controls for both technical and biological factors and highlights complex cellular relations and biological insights.

## 1 Introduction

Single cell genomics – especially single cell RNA-seq (scRNA-seq) – has opened the way to comprehensive analysis of the relationship between cells, including their different types, states, physiological transitions, differentiation trajectories, and spatial positions (Regev et al., 2017; Stegle et al., 2015; Wagner et al., 2016). Although scRNA-seq datasets have high dimensionality, their intrinsic dimensionality is typically low, because many genes are co-expressed and a few variables, such as cell type, a gene program, or the number of detected transcripts, could explain a substantial portion of the variation in a dataset. As a result, dimensionality reduction, followed by visualization or downstream analyses has become a key strategy for exploratory data analysis in single cell genomics (Butler et al., 2018; Luecken and Theis, 2019).

Recently, deep learning models (Ching et al., 2018), especially (variational) auto-encoders (Kingma and Welling, 2013; Kingma et al., 2014; Rezende et al., 2014), have been used for dimensionality reduction prior to visualization or downstream analyses, such as clustering (Ding et al., 2018; Eraslan et al., 2019; Grønbech et al., 2019; Lopez et al., 2018; Lotfollahi et al., 2019; Wang and Gu, 2018). This leverages their ability to model large-scale high-dimensional data and their flexibility in incorporating different factors, especially batch effects in the modeling framework.

However, standard variational auto-encoders have several shortcomings when modeling and analyzing scRNA-seq data. First, they assume a standard multi-dimensional normal prior for the low-dimensional latent variables. Unfortunately, this prior encourages the low-dimensional representations of all cells to group in the center of the latent space, even for data consisting of distinct cell types, especially if the model is trained long enough, such that the posterior distributions gradually approximate the prior distribution. A second challenge arises from using the cosine to measure the distance between two cells (Bendall et al., 2014; Haghverdi et al., 2018; Kiselev et al., 2018) for very sparse droplet-based scRNA-seq data (>90% genes with zero counts in a typical cell profile). Because the cosine distance between two cell vectors is their Euclidean distance after normalizing the two cell vectors to have a unit *ℓ*^2^ norm, the cells lie on the surface of a unit hypersphere with a dimensionality of *D* − 1, where *D* is the number of measured genes. Embedding data distributed on a hypersphere to a Euclidean space introduces significant distortion for commonly-used dimensionality reduction tools (Cooley et al., 2019), and standard variational auto-encoders also fail to model such data (Davidson et al., 2018). Finally, the Euclidean geometry is not optimal for representing hierarchical, branched developmental trajectories (Klimovskaia et al., 2019; Nagano et al., 2019; Nickel and Kiela, 2018).

Here, we present new approaches for embedding of cells into hyperspherical or hyperbolic spaces, to better capture their inherent properties. For general scRNA-seq data, we minimize the distortion by embedding cells to a lower-dimensional hypersphere instead of a low-dimensional Euclidean space (Davidson et al., 2018), using von Mises-Fisher (vMF) distributions on hyperspheres as the posteriors for the latent variables (Davidson et al., 2018; Guu et al., 2018; Xu and Durrett, 2018). Because the prior is a uniform distribution on a unit hypersphere and the uniform distribution on a hypersphere has no centers, points are no longer forced to cluster in the center of the latent space. For representation and inference of hierarchical, branched developmental trajectories, we embed cells to the hyperbolic space of the Lorentz model and visualize the embedding in a Poincaré disk (Mathieu et al., 2019; Nagano et al., 2019; Nickel and Kiela, 2018). As we show across six diverse datasets, our model results in enhanced visualization, while simultaneously addressing complex batch effects, thus providing an important tool for single cell genomics research.

## 2 Results

### 2.1 Mapping scRNA-seq data to hyperspherical or hyperbolic latent spaces

We developed scPhere (pronounced “sphere”), a deep learning method that takes scRNA-seq count data and information about multiple known confounding factors (*e.g.*, batch, condition) and embeds the cells to a hyperspherical or hyperbolic latent space (**Fig. 1A, Methods**). We reasoned that scPhere would allow cells to be embedded more appropriately, because they will not be constrained to aggregate in the center. In cases where we expect a branching or hierarchical structure, hyperbolic spaces are particularly suitable, because the exponential volume growth of hyperbolic spaces with radius confers them enough capacity to embed trees, which have exponentially increasing numbers of nodes with depth. For 3D visualization, scPhere places cells on the surface area of a sphere (but not inside the sphere), such that we only need to rotate the sphere to see all cells. The scPhere package renders all 3D plots for interactive visualizations of millions of cells with the rapid rgl R package, with web graphics library files, which can be opened in a browser for exploration.

**Figure 1.**
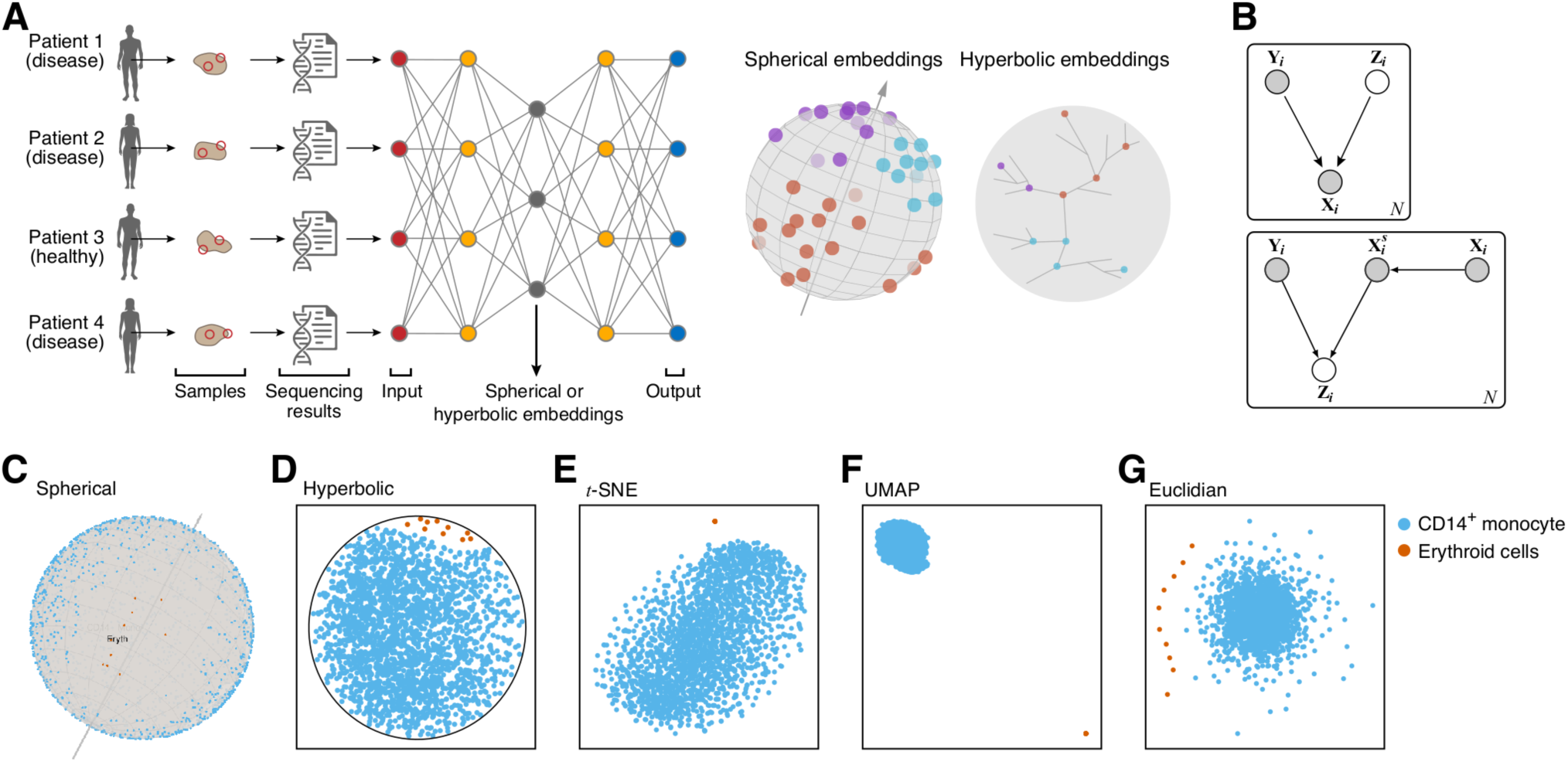
scPhere model. (**A**) Method overview. (**B**) scPhere directed probabilistic graphical model and the variational approximation of its posterior. ***Y***_*i*_ is the categorical batch variable of a cell *i*, ***Z***_*i*_ is the low-dimensional latent variable for the cell distributed on a hypersphere (or in a hyperbolic space), and ***X***_*i*_ is the raw UMI count vector of cell *i*. For the variational approximation of the posterior of ***Z***_*i*_, ***X***_*i*_ is first *log*-transformed and scaled to have a unit *ℓ*^2^ norm (represented by 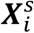) if a hyperspherical latent space is used, otherwise ***X***_*i*_ is only *log*-transformed without scaling. (**C-G**) scPhere addresses the crowding challenge compared to other methods. Shown are different embedding of single cell profiles (dots) color coded by type, (**c**) scPhere posterior mean (a dot on the surface of the unit sphere); (**D**) scPhere learned representation in the Poincaré disk; (**E**) a 2D *t*-stochastic neighborhood embedding (*t*-SNE), (**F**) a 2D uniform manifold approximation and projection (UMAP) representation, and (**G**) a 2D latent representation from scPhere with a standard normal prior and normal posterior.

Specifically, scPhere takes as input an scRNA-seq dataset 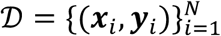 with *N* cells, where ***x***_*i*_ is the UMI count vector of cell *i*, and ***y***_*i*_ is a category vector specifying the batch in which ***x***_*i*_ is measured, and models the ***x***_*i*_ UMI count distribution as governed by a latent low-dimensional random vector ***z***_*i*_ and by ***y***_*i*_ (**Fig.** 1**B**). Note, that ***y***_*i*_ can account for multiple confounding factors, for example, the patient, disease status, and lab protocol. The scPhere model assumes that the latent low-dimensional random vector ***z***_*i*_ is distributed according to a prior, with the joint distribution of the whole model factored as *p*(***y***_*i*_ | ***θ***_*i*_)*p*(***z***_*i*_ | ***θ***_*i*_)*p*(***x***_*i*_ | ***y***_*i*_, ***z***_*i*_, ***θ***_*i*_), where *p*(***y***_*i*_ | ***θ***_*i*_) is the categorical probability mass function (constant for our case, as ***y***_*i*_ is observed). For hyperspherical latent spaces, scPhere uses a uniform prior on a hypersphere for *p*(***z***_*i*_ | ***θ***_*i*_); for hyperbolic latent spaces, it uses a wrapped normal distribution in the hyperbolic space as the prior. For the observed raw UMI count inputs, we assume a negative binomial distribution: 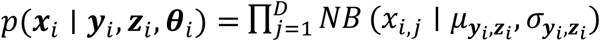, with parameters specified by a neural network. The inference problem is to compute the posterior distribution *p*(***z***_*i*_ | ***y***_*i*_, ***x***_*i*_, ***θ***_*i*_), which is assumed to be a von-Mises-Fisher distribution for hyperspherical latent spaces, and a wrapped normal distribution for hyperbolic latent spaces. Because it is intractable to compute the posterior, the scPhere model uses a variational distribution *q*(***z***_*i*_ | ***y***_*i*_, ***x***_*i*_, ***ϕ***_*i*_) to approximate the posterior (**Fig. 1B**). When a hyperspherical latent space is used, ***x***_*i*_ is first log-transformed and scaled to have a unit *ℓ*^2^ norm for inference, otherwise ***x***_*i*_ is only log-transformed but not scaled. The parameters ***ϕ***_*i*_ of the variational distribution are (continuous) functions of ***x***_*i*_ and ***y***_*i*_ parameterized by a neural network with parameter ***ϕ***. We provide full details in the **Methods** section.

### 2.2 Spherical latent variables help resolve cell crowding

Applying scPhere to scRNA-seq data shows that its spherical latent variables help address the problem of cell crowding in the center, and that it provides an excellent visualization for data exploration, with latent variable posterior means of cells which are easily interpretable. As a first illustrative test, when we applied scPhere with a hyperspherical latent space to a data set (Stoeckius et al., 2017) with only 10 erythroid cell profiles and 2,293 CD14^+^ monocytes, the 10 erythroid cells were close to each other and relatively far from the CD14^+^ monocytes (**Fig. 1C**). Moreover, the posterior means of cells typically did not overlap, which helped ensure that if cells are colored by gene expression, we can discern individual cells without occlusion. When we used a hyperbolic (instead of hyperspherical) latent space, the erythroid cells were still close to each other and far from the center (**Fig. 1D**). By comparison, both *t-*stochastic neighborhood embedding (*t*-SNE) (Maaten and Hinton, 2008) and uniform manifold approximation and projection (UMAP) (McInnes et al., 2018) put CD14^+^ monocytes at the origin (**Fig. 1E,F**), and the 10 erythroid cells were embedded so close to each other that they almost collapsed to a single point. Finally, when we used a standard multivariate normal prior, the posterior means of the latent variables were centered at the origin, leading to crowding of the CD14^+^ monocytes and peripheral spreading of the 10 erythroid cells (**Fig. 1G**).

This result generalized across multiple datasets of diverse biological systems, including 3,314 human lung cells (Braga et al., 2019) (**Fig. 2A**), 1,378 mouse white adipose tissue stromal cells (Hepler et al., 2018) (**Fig. 2B**), and 1,755 human splenic nature killer cells spanning four subtypes (Crinier et al., 2018) (**Fig. 2C**): in each case, cells of the same type were close to each other on the surface of a sphere, and yet generally two cells are distinguishable, even by eye (**Fig. 2A-C**). Conversely, in the Euclidean space, the closer the cells were to the center, the higher were their densities (**Fig. 2D-F**), a problem persisting in both 2D (**Fig. 2D-F**) and 3D (**Supplementary Fig. 1A-C**), even with rotation of the 3D space. In particular, similar cell types were very close to each other in the Euclidean space (*e.g.*, APC and FIP, **Fig. 2E**), and rare cell types became ‘outliers’ (hNK_Sp3 and hNK_Sp4, **Fig. 2E**).

**Figure 2.**
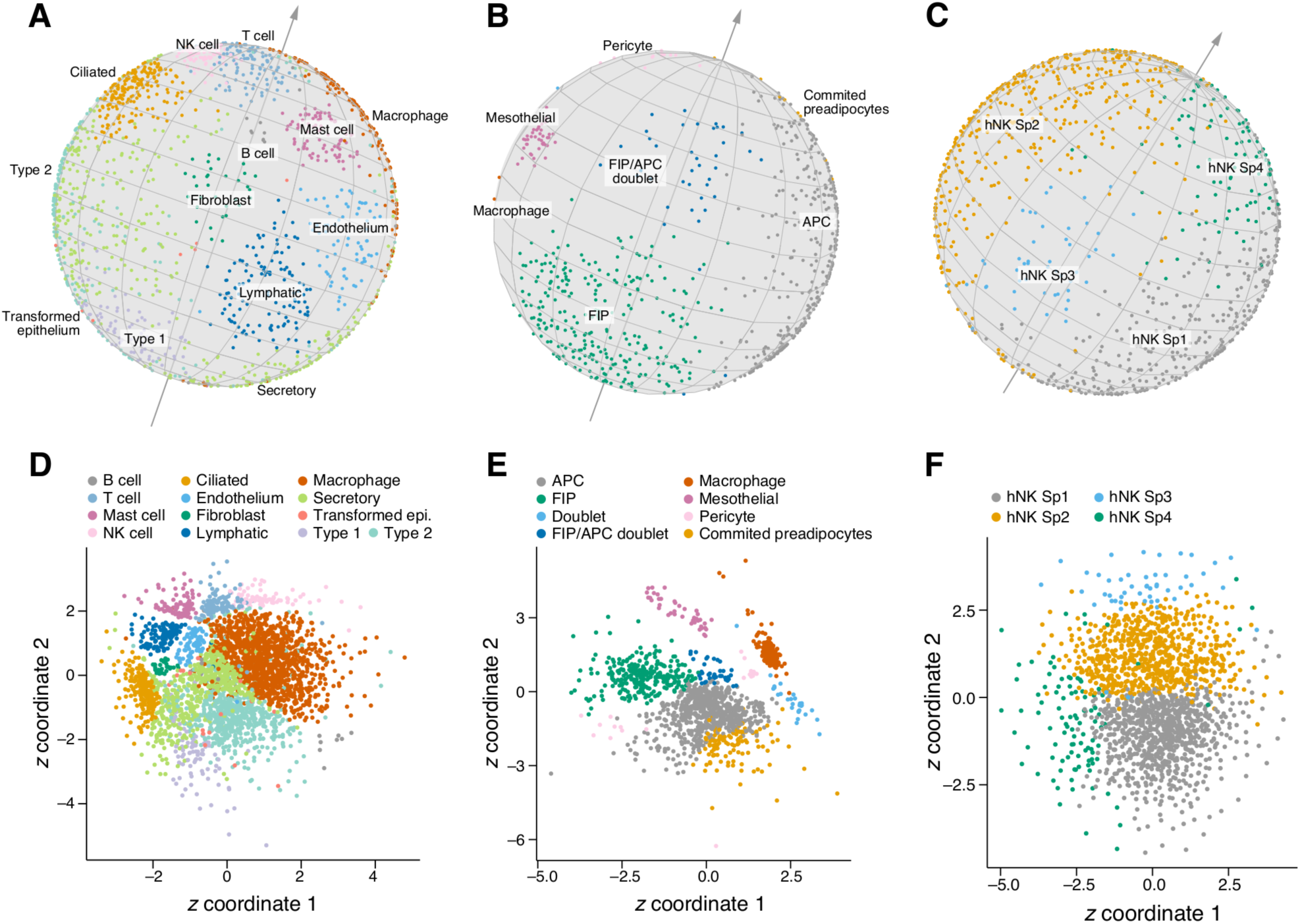
scPhere addresses the cell crowding problem by using a spherical latent space. scPhere embeddings on the surface of unit spheres (**A-C**) and in the Euclidean space (**D-F**) of 3,314 human lung cells (**A,C**), 1,378 mouse white adipose tissue stromal cells (**B,E**), and 1,755 human splenic natural killer cells (**C,F**), color coded by their annotated types.

### 2.3 Spherical latent variables effectively model batch and other variables

ScPhere gracefully addresses batch correction, which we illustrate by its application to 301,749 cells we previously profiled in a complex experimental design from the colon mucosa of 18 ulcerative colitis (UC) patients and 12 healthy individuals (Smillie et al., 2019). These cells were collected separately from the epithelial and lamina propria fractions of each biopsy, in two replicate biopsies for each healthy individual and as a pair of inflamed and uninflamed biopsies for the UC patients (for a few UC patients, there were replicate inflamed and/or replicate uninflamed biopsies). Moreover, samples were collected at two time periods, separated by over a year (and analyzed as “train” and “test” data in the original study (Smillie et al., 2019)).

Analyzing the stromal and glia cells, taking patient origin of cells as the batch vector, not only recapitulated the main cell groups in our initial study (Smillie et al., 2019), but was highly refined, allowing us to better explore cellular relations (**Fig. 3A, Supplementary Movie 1**). For example, endothelial cells and microvascular cells were close to each other, and adjacent to post-capillary venules. Conversely, these distinctions can barely be discerned in a UMAP plot of the same data (**Fig. 3B**; using the 20 batch-corrected principal components by Harmony (Korsunsky et al., 2018) as inputs). Among fibroblasts, cells arranged in a manner that mirrored their position along the crypt-villus axis, from *RSPO3*^+^*WNT2B*^+^ cells (which likely support the ISC niche (Smillie et al., 2019)), to *WNT2B*^+^ cells, and to *WNT5B*^+^ cells. Strikingly, the inflammatory fibroblasts were readily visible (**Fig. 3A**, light blue), and were both distinctive from the other fibroblasts, while spanning the range of the “crypt-villus axis” (as shown experimentally (Smillie et al., 2019)). To demonstrate scPhere’s ability to correct for multiple confounding factors, we reran it using both patient origin and disease status (healthy, uninflamed, inflamed) as the batch vector. This merged the inflammatory fibroblasts with *WNT2B*^+^ fibroblasts (**Fig. 3C, Supplementary Movie 2**), as the influence of disease status on cell types was largely removed, which may suggest their origin.

**Figure 3.**
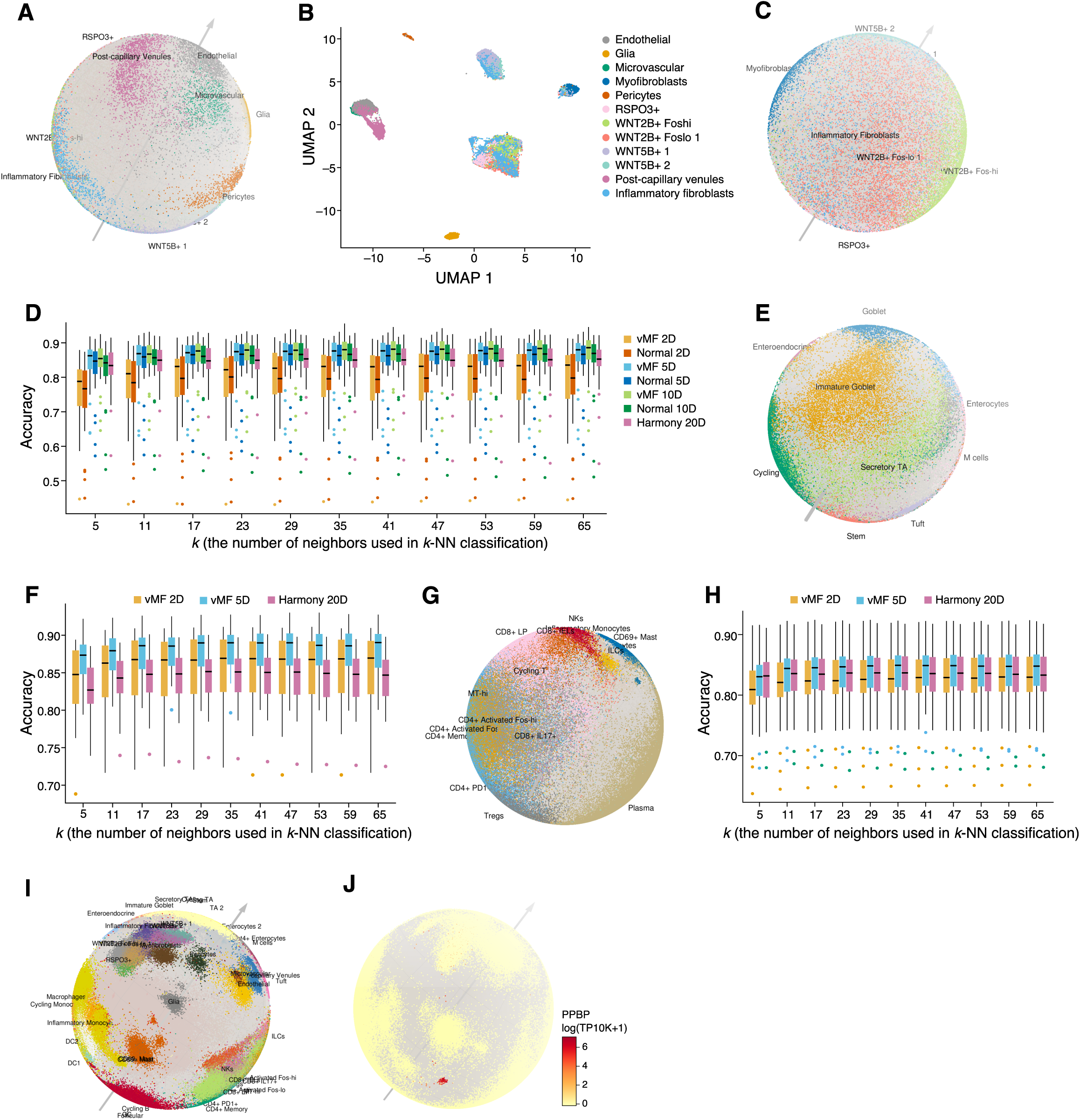
Effective visualization, batch correction and classification of a colon mucosa atlas with scPhere. **(A-C)** scPhere correct for patient and disease status in embedding of stroma cells. Embedding of stromal cells and glia cells on a sphere, with patient origin of cells as the batch vector **(A)**, by UMAP embedding with 20 batch-corrected principal components from Harmony as inputs **(B)**, or on a sphere when both the patient and disease status were used as the batch vector **(C). (D,F,H)** Successful classification in low dimensions. *k*-nearest neighbor classification accuracy of stromal **(D)**, epithelial **(F)** or immune **(H)** cell types (y axis), for different *k*’s (x axis), when testing on the cells from one patient, after training on the cells from all other patients, with each of several methods (color legend). If a normal prior and normal posteriors were used, gene expression count vectors were only log transformed without normalizing to have unit *ℓ*^2^ norm for inference. (**E,G,I**) ScPhere embedding of epithelial (**E**), immune (**G**) and all 300,000 stromal, epithelial, and immune cells simultaneously (**I**). (**J**) ScPhere highlights rare cell subsets. A small subset of megakaryocytes/platelets, colored by PPBP expression level (color bar, log(transcripts per 10,000) or log(TP10K+1)).

Batch vectors help highlight the influence of both disease and anatomical regions on gene expression from scPhere’s latent representations. For example, in analyzing epithelial cells, which are derived from both the epithelial and lamina propria (LP) samples, when the anatomical regions were not used as a component of the batch vector, some of the cells from the LP and epithelial layer organized in two respective parallel tracts in some regions of the sphere (**Supplementary Fig. 2A**, orange *vs*. grey), with each tract including both stem cells, Transit Amplifying 2 (TA2) cells, cycling TA cells, and immature enterocytes (**Supplementary Fig. 2B**). However, when we added anatomical regions as a component of the batch vector, the cells were grouped solely by types (*e.g.*, stem cells separate from TA2 cells) instead of region (**Supplementary Fig. 2C,D**). Notably, cell types that were mostly from one region (*e.g.*, tuft cells, mostly from epithelial fractions) remained grouped distinctly (**Supplementary Fig. 2C,D**). Similarly, when we did not use disease status (healthy, uninflamed, or inflamed) as a component of the batch vector, some cells (*e.g.*, TA2, immature enterocytes, and enterocytes) had some ‘outliers’ mapped to low-density regions of the sphere (**Supplementary Fig. 2E**), mostly from UC (uninflamed or inflamed) samples (**Supplementary Fig. 2F**). Once we also used the disease status of each cell as a component of the batch vector, these cells formed more compact clusters (**Supplementary Fig. 2G**), and the cells of different disease states (**Supplementary Fig. 2H**) and from different patients (**Supplementary Fig. 2I**) were well mixed. In this manner, a user can understand cellular relations driven by different biological factors.

### 2.4 scPhere preserves the structure of scRNA-seq data even in very low-dimensional spaces

For embedding in a latent space with few dimensions, scPhere introduced fewer distortions in the dimensionality-reduction process compared to the case with a latent Euclidean space, as reflected by performance on tasks such as cell classification. To assess this, we first performed dimensionality reduction on the colon dataset and then trained a *k*-nearest neighbor (*k*-NN) classifier on the cells (using the labels from the original study (Smillie et al., 2019)), holding out cells from one patient at a time for testing. We compared the classification accuracy obtained using scPhere’s hypersphere embedding to that obtained when we run it using a standard normal prior and normal posteriors, which embeds cells in a Euclidean latent space. When using only two dimensions (**Fig. 3D**), the results from scPhere were significantly better than when using a Euclidean latent space across all *k*s (FDR < 0.05, paired *t*-test, two-tailed), suggesting that compared to a Euclidean latent space, a hyperspherical latent space introduced less distortions when the latent dimensionality was low. This lesser distortion in low-dimensions is especially useful for data visualizations, because these typically use a two-dimensional latent space. As expected, *k*-NN classification accuracies increased for both choices with the number of latent dimensions, suggesting less distortions introduced by dimensionality reduction (**Fig. 3D**). ScPhere’s classification results were also comparable with those from Harmony (Korsunsky et al., 2018) when the latter used 20 principal components (**Fig. 3D**).

### 2.5 The preserved structure aids in classification and visualization

As a parametric model, we can train scPhere to co-embed unseen (test) data to a latent space learned from training data only. To demonstrate this, we performed a 10-fold cross-validation analysis, where we partitioned the colon fibroblasts and glia cells into 10 roughly equally-sized subsamples, held out one subsample as out-of-sample evaluation data, and used the remaining nine subsamples as training data to select variable genes and learn different scPhere models to embed cells on a 5D hypersphere. We then trained a *k*-NN classifier on the 5D representations of the training data and used the *k*-NN classifier to classify the 5D representations of the out-of-sample evaluation data. We repeated this process 10 times with each of the 10 subsamples used exactly once as the out-of-sample validation data. The *k*-NN classifiers had a median accuracy of 0.834-0.853 (*k* = 5 or 65, respectively, **Supplementary Fig. 3**). By comparison, when we repeat this process but using pre-computed 5D representations from all fibroblasts and glia cells, accuracy was similar (0.847-0.860, the minimal two-tailed Wilcoxon signed-rank test FDR=0.036, and for two *k*s, the FDRs > 0.05, **Supplementary Fig. 3**).

This good performance in visualization, batch correction and classification was also apparent for the epithelial and immune cells, where we included patient origin, disease status, and anatomical region as the batch vectors. For epithelial cells, cells grouped visually by type (rather than other variables), and ordered in a manner consistent with their development: stem cells → secretory Transit Amplifying (TA) cells → immature goblet cells (or cycling TA) → goblet cells (Smillie et al., 2019) and stem cells → TA2 cells → immature enterocytes → enterocytes (**Fig. 3E, Supplementary Movie 3**). Cell classification was highly accurate (ranging from 0.87 for *k* = 5, to 0.89 for *k* = 65, **Fig. 3F**), and the influence of region, disease status, and patient was largely removed (**Supplementary Fig. 2D,H,I**). Notably, scPhere classification accuracy on a 5D latent space significantly outperformed Harmony with 20 principal components (FDR < 10^−6^ for all *k*s, paired *t*-test, two-tailed). In immune cells, the three major cell classes (B cells, T cells, and myeloid cells, **Fig. 3G**) were readily identifiable, and well organized relative to each other. In particular, CD8^+^IL17^+^ T cells were nestled between CD8^+^ T cells and activated CD4^+^ T cells in a manner that was intriguing and consistent with the mixed features of those cells (Smillie et al., 2019), which are CD8 T cells that also express low levels of CD4 protein (Smillie et al., 2019) and an *IL17* module (Smillie et al., 2019) typically associated with CD4^+^ T cells. Notably, the *k*-NN classification accuracies were lower in immune than in epithelial cells (**Fig. 3H**), reflecting mostly the continuum of T cell states, which is less well captured as disjoint classes, but similar using a 2D latent space, 5D latent space, or Harmony (**Fig. 3H**). Finally, when we analyzed the immune, stromal cells and epithelial cells simultaneously, the results were quite similar to those from analyzing these cells separately (**Fig. 3I, Supplementary Movie 4**), demonstrating the capacity of scPhere to embed large numbers of cells of diverse types, states and proportions.

### 2.6 Clustering cells following scPhere embeddings

To demonstrate how scPhere impacts clustering analysis, we clustered (using the Louvain algorithm (Blondel et al., 2008; Levine et al., 2015)) the embeddings of cells on the surface of 5D hyperspheres and compared them to the clusters in the original study (Smillie et al., 2019) (where only patients were used as the batch vector and variable genes were selected for each patient separately to compute a census of batch-insensitive variable genes (Smillie et al., 2019)). The fibroblasts and glia cells were partitioned into 18 clusters (**Supplementary Fig. 4A**), which were largely consistent with the original analysis (Smillie et al., 2019) with some minor exceptions: *RSPO3*^+^ fibroblasts included cells from the original *WNT2B*^+^ Fos-lo cluster, and some of the inflammatory fibroblasts were in the *WNT2B*^+^ fibroblast clusters, highlighting their molecular similarity. We obtained similar clustering results when we used cell embeddings on the surface of a 10D hypersphere (**Supplementary Fig. 4B**), consistent with our classification results (**Fig. 3D**). Clustering the epithelial cells in the 5D hyperspherical space (we included patient origin, disease status, and anatomical region as the batch vectors) produced 20 clusters (**Supplementary Fig. 4C**): 12 map one-to-one to the previous 12 annotated cell subsets, whereas some of the larger subsets were split further (cluster 1, 3, 7). Some clusters consisted of cells from multiple subsets. For example, cluster 12 (from secretory TA: cluster 10, TA2: cluster 0, and immature enterocytes: cluster 4) cells co-expressed TA2 marker genes (*e.g., CA2*) and secretory TA marker genes (*e.g*., *ZG16*). These may reflect a transition state in the epithelial differentiation continuum, but may also be cell doublets; further studies may help distinguish these possibilities. Clustering the immune cells on the surface of a 5D hypersphere produced 35 clusters (**Supplementary Fig. 4D, Supplementary Movie 5**). Some cells with very similar molecular features were merged (CD4^+^ Activated Fos-hi and CD4^+^ Activated Fos-low, **Supplementary Fig. 4D**). Some of these merged cells were differently distributed in regions (CD69^-^ mast cells and CD69^+^ mast cells between Epithelial and LP regions; **Supplementary Fig. 4E**), or disease (cycling monocytes and macrophages between healthy, uninflamed, or inflamed, **Supplementary Fig. 4F**) as we corrected for the influences of region, disease, and patient.

Notably, rare cell types were also distinct in the low-dimensional space. For example, a small cluster consisted of megakaryocytes/platelets (**Fig. 3J, Supplementary Fig. 4E, cluster 33**), expressing *PPBP, PF4, GNG11, NRGN, TUBB1*, that were missed in the original analysis. Other examples included a small B cell cluster (cluster 34) exclusively expressing *IGLC7* and a small monocyte cluster (cluster 28) expressing *FCGR3A* and *RHOC*.

### 2.7 Embedding cells in a hyperbolic space for trajectory discovery and interpretation

When cells are expected to show developmental trajectories, such as from adult stem cells to differentiated cells, scPhere can embed them into a hyperbolic space of the Lorentz model (Nagano et al., 2019; Nickel and Kiela, 2018), and optionally convert the coordinates in the Lorentz model to the Poincaré disk for visualization (Mathieu et al., 2019; Nickel and Kiela, 2017).

Applying this first to epithelial cells, we readily discerned developmental ordering from intestinal stem cells to terminally differentiated cells in either the Poincaré disk (**Fig. 4A**), or in the Lorentz model (**Supplementary Fig. 5A**): the two major cell development trajectories are clearly delineated and M-cells and Best^+^ enterocytes are close to each other. PHATE (Moon et al., 2019) (multidimensional scaling on top of diffusion maps) analysis using the 5D representations of cells in the Lorentz model as inputs recapitulated the results from the 2D representations (**Supplementary Fig. 5B**). In contrast, developmental trajectories were less apparent when we embedded cells in a Euclidean space (**Fig. 4B**), with the two major differentiation trajectories located <u>closely</u> on one side of the 2D plane, or when we used PHATE multidimensional scaling on the 5D representations of cells in the Euclidean space (**Supplementary Fig. 5C**), where cells in the two major developmental branches were again close to each other.

**Figure 4.**
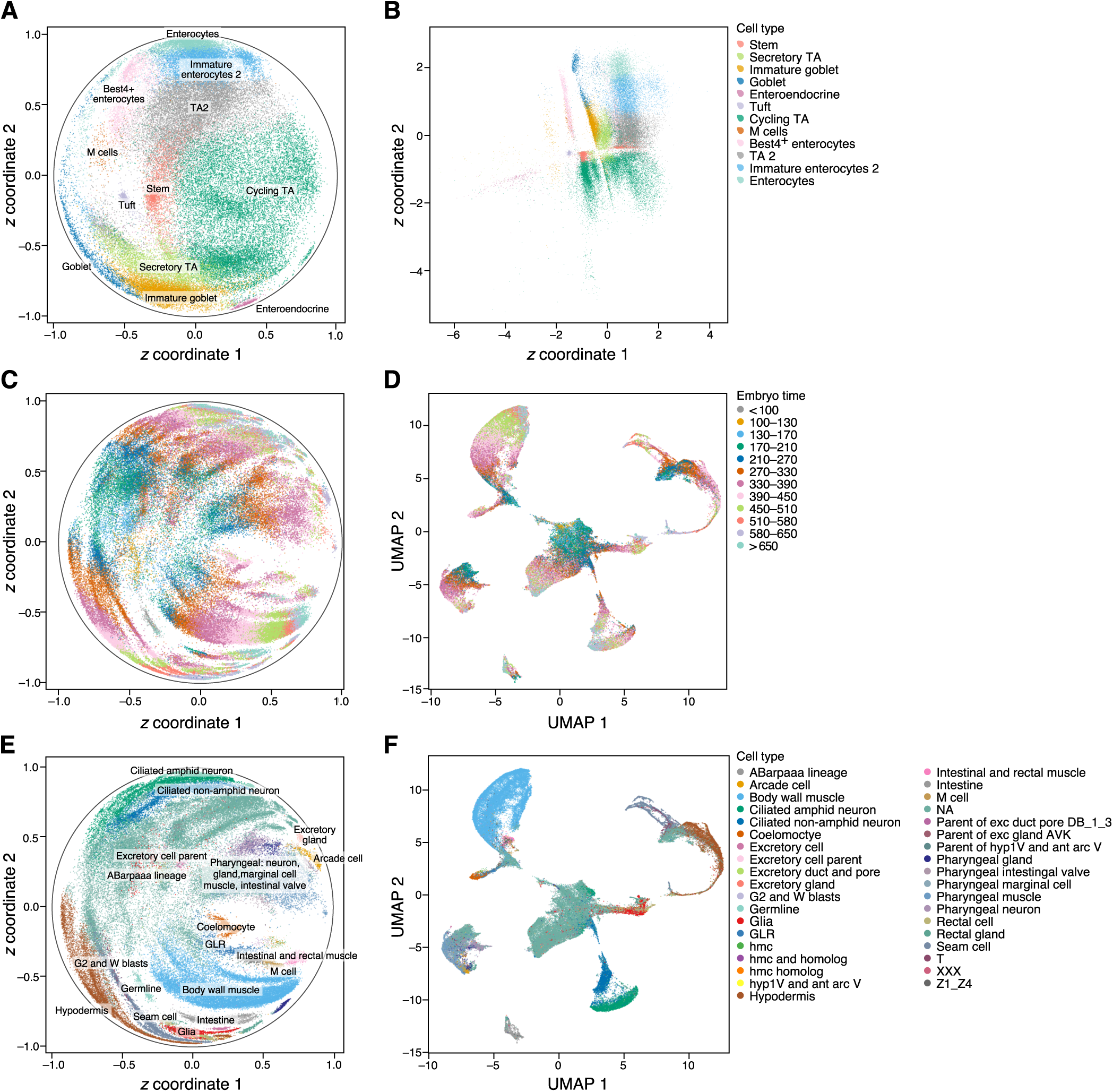
Embedding cells in the hyperbolic space for trajectory discovery and interpretation. **(A,B)** Major development trajectories of colon epithelial cells are discerned from embeddings in the Poincaré disk. Single epithelial cell profiles (dots) annotated by type as embedded in a Poincaré disk **(A)** or the Euclidean space **(B). (C-F)** Embedding of cells from a *C. elegans* embryonic time course highlights both developmental timing and differentiation to subsets. Single epithelial cell profiles (dots) annotated by either time (**C,D)** or cell type **(E,F)** in a hyperbolic space of the Poincaré disk **(C,E)** or in a 2D UMAP (with 20 principal components, batch corrected by Harmony) **(D,F).**

Finally, when we analyzed 86,024 *C. elegans* embryonic cells (Packer et al., 2019) collected along a time course from <100 minutes to >650 minutes after each embryo’s first cleavage, cells were ordered neatly in the latent space by both time and lineage, from a clearly discernible root at time <100 to cells from time >650 near the border of the Poincare disk **(Fig. 4C, Supplementary Fig. 6**) or away from the origin in the Lorentz model (**Supplementary Fig. 7A**). These patterns are harder to discern in a UMAP (**Fig. 4D**, with 20 batch-corrected PCs by Harmony as inputs). Importantly, within the same cell type, the cells were ordered by embryo time in the Poincaré disk (**Fig. 4E**) or in the Lorentz model (**Supplementary Fig. 7B**). For example, the cells of the body wall muscle (BWM, as annotated in (Packer et al., 2019), the most abundant cell type in this dataset, **Supplementary Fig. 6**) first appeared at embryo time 130-170 in a separable position, and then “advance” towards the disc’s periphery in a continuous progression but in a manner aligned with embryo time (*i.e.*, from 170 - 210 to >650) and lineages (i.e., from first row and second row BWMs (D and MS lineage) to posterior BWMs (C lineage) (Packer et al., 2019), **Supplementary Fig. 8**). Conversely, similar cell types were visually indistinguishable in the UMAP (**Fig. 4F**). Thus, the scPhere model with a hyperbolic latent space help represent developmental and other temporal processes.

## 3 Discussion

We introduced scPhere, a deep generative model to embed single cells on hyperspheres or in hyperbolic spaces. When a cosine distance is used to measure the distance between two cells, a spherical latent space is the natural choice for embedding scRNA-seq data. By using a hyperspherical latent space, scPhere overcomes the problem of forcing cells to center at the origin of the latent space. This provides more readily interpretable representations, and avoids occlusion, as we demonstrate in diverse systems, including >300,000 epithelial, immune and stromal cells from the colon mucosa. Moreover, by embedding cells in hyperbolic spaces, scPhere helps discovering and interpreting developmental trajectories, as we show for both asynchronous differentiation of epithelial stem cells and for 86,024 cells collected along a time course of *C. elegans* embryonic development. ScPhere thus enhances future exploratory data analysis and visualization of cells from single-cell studies.

scPhere also effectively accounts for multiple complex batch effects, which we show disentangles cell types from patients, diseases, and location variables, and facilitates downstream analyses. As batch correction is generally a challenging task, and in the future, we can leverage supervised information to further provide uncertainties of aligning cells from batches.

The scPhere model is robust to hyper-parameters. Here, we used the same hyper-parameters for scPhere analyses for all datasets (the number of cells varies across datasets, from ∼1,000 to >300,000), whereas some previous studies (Hu and Greene, 2019) showed that classical variational autoencoders could be sensitive to hyper-parameters. ScPhere’s robustness may stem from the robust negative binomial distribution for modeling UMI counts, or from the use of non-Euclidean latent spaces to help solve the cell-crowding problem in the latent space.

ScPhere can be extended in several ways. When cell type annotations or cell type marker genes for some of the analyzed cells are available, we can include semi-supervised learning to annotate cell types (Xu et al., 2019; Zhang et al., 2019). Given the rapid development of spatial transcriptomics (Rodriques et al., 2019; Vickovic et al., 2019), single-cell ATAC-seq (Lareau et al., 2019; Satpathy et al., 2019) and other complementary measurements, scPhere can be extended for integrative analysis of multi-modality data. We can also learn discrete hierarchical trees for better interpreting developmental trajectories. Given the large number of cells to be sequenced by large international initiatives such as the Human Cell Atlas (Regev et al., 2017), we foreseen that scPhere will be a valuable tool for large-scale single-cell genomics studies.

## 4 Methods

### 4.1 Mapping scRNA-seq data to a hyperspherical latent space

ScPhere received as input a scRNA-Seq dataset 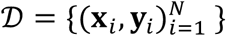, where **x**_*i*_ ∈ ℝ^*D*^ is the gene expression vector of cell *i, D* is the number of measured genes, **y**_*i*_ is a categorical variable vector specifying the batch in which **x**_*i*_ is measured, and *N* is the number of cells. Although **x**_*i*_ is high-dimensional, its intrinsic dimensionality is typically much lower. We therefore assume that the **x**_*i*_ distribution is governed by a much lower-dimensional vector **z**_*i*_, and the joint distribution is factorized as follows (**Fig. 1b**):

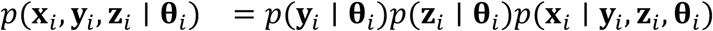

Here *p*(**y**_*i*_ | **θ**_*i*_) is the categorical distribution, *p*(**z**_*i*_ | **θ**_*i*_) is the prior distribution for **z**_*i*_ (**z**_*i*_ ∈ ℝ^*M*^, **z**^*T*^**z** = 1, *M* ≪ *D*), which is assumed to be a uniform distribution on a hypersphere with density 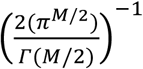. For notational simplicity, we use bold font **θ**_*i*_ to represent the parameters of each distribution, *e.g.*, the parameters **θ**_*i*_ in *p*(**y**_*i*_ | **θ**_*i*_) and *p*(**z**_*i*_ | **θ**_*i*_) are the parameters of the two distributions and should be different.

For scRNA-seq data, the observed Unique Molecular Identifier (UMI) count of gene *j* in cell *i* has typically been assumed to follow a zero inflated negative binomial (ZINB) distribution (Eraslan et al., 2019; Lopez et al., 2018; Pierson and Yau, 2015). However, a recent study suggests that zero inflation is an artifact of normalizing UMI counts (Townes et al., 2019), and negative binomial distributions generally fit the UMI counts well (Hafemeister and Satija, 2019; Svensson, 2019; Vieth et al., 2017). We therefore assume a negative binomial distribution of observations in this study:

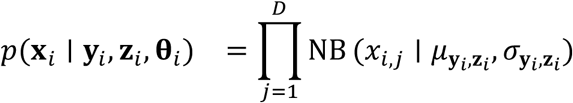

The negative binomial parameters mean 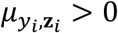 and dispersion 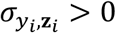 are specified by a model neural network (decoder), which can model complex nonlinear relationships between the latent variables and the observations.

We next want to compute the posterior distribution *p*(**z**_*i*_ | **y**_*i*_, **x**_*i*_, **θ**_*i*_), which is assumed to be a von-Mises-Fisher distribution on a unit hypersphere of dimensionality *M* − 1: 𝕊^*M*−1^ = {**z** | **z** ∈ ℝ^*M*^, **z**^*T*^**z** = 1}. We turn to variational inference to find a *q*(**z**_*i*_ | **y**_*i*_, **x**_*i*_, **ϕ**_*i*_) to approximate the posterior, since exact inference is intractable, given that the model is parameterized by a neural network. In addition, the number of parameters to estimate grows with the number of cells, because each cell has a ‘local’ distribution with parameter **ϕ**_*i*_. To scale to large datasets, variational auto-encoders use an inference neural network (encoder, with a fixed number of parameters) to output the ‘local’ parameter **ϕ**_*i*_ of each cell. Therefore, the learning objective is to find the model neural network and the inference neural network parameters to maximize the evidence lower bounds:

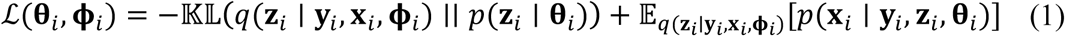

The Kullback-Leibler (𝕂𝕃) divergence (Kullback and Leibler, 1951) in **Equation 1** can be calculated analytically (below). We use Monte-Carlo integration (sampling from the vMF distribution *q*(**z**_*i*_ | **y**_*i*_, **x**_*i*_, **ϕ**_*i*_)) to calculate the second term.

To make scPhere robust to small perturbations (*e.g.*, sequencing depth), we add a penalty term to the objective function in **Equation 1**. Specifically, for each gene expression vector **x**_*i*_, we down-sample **x**_*i*_ by keeping 80% of its UMIs to produce 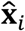. The latent representations of **x**_*i*_ and 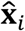 are **z**_*i*_ and 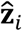, respectively. The penalty term is defined as 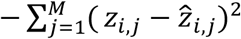 as we want **z**_*i*_ and 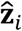 to be close.

### 4.2 von-Mises-Fisher distribution

The von-Mises-Fisher (vMF) (Mardia and El-Atoum, 1976) represents angular observations as points on the surface of a unit-radius hypersphere. Let **z** be a *M*-dimensional random vector with unit radius (**z**^*T*^**z** = 1), then its probability density function is:

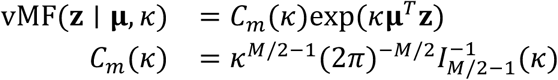

where **μ**^*T*^**μ** = 1 is the mean direction vector (not the mean) and *κ* ≥ 0 is the concentration parameter. The greater the value of *κ*, the higher the concentration of distribution around the mean direction vector **μ**. When *κ* = 0, 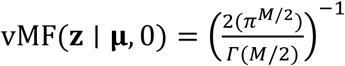 is the uniform distribution on the unit hypersphere 𝕊^*M*−1^. *C*_*m*_ (*κ*) is a constant normalization factor and *I*_*ν*_(·) is the modified Bessel function of the first kind of order _*ν*_ (Straub et al., 2015): 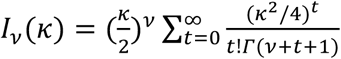. The Gamma function is defined as 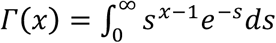.

### 4.3 The Kullback-Leibler divergence between the posterior and prior

For random vectors distributed on the surface of a hypersphere, a natural prior is the uniform distribution, which is the vMF distribution with zero concentration vMF(_**Z**_ | **μ**, 0). In this case, the Kullback-Leibler (𝕂𝕃) divergence (Kullback and Leibler, 1951) can be written in closed-form:

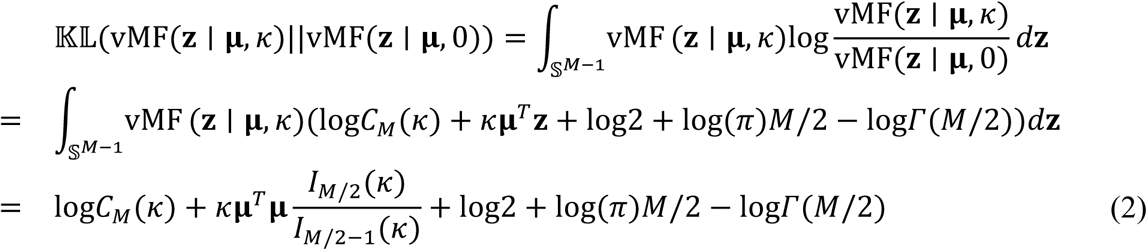

with

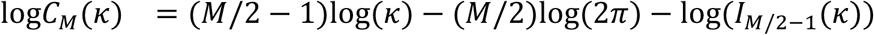

Notice that **Equation 2** is independent of the mean direction vector **μ** as **μ**^*T*^**μ** = 1, so we only need to take the derivative of **Equation 2** w.r.t *κ* during optimization. In other words, minimizing the 𝕂𝕃 divergence only forces the concentration parameter *κ* to be close to zero but without any forces on the mean direction vector. This is different from using a location-scale family of priors, such as a standard normal prior, where the prior encourages the posterior means of all points to be close to zero. When *ν* ≪ *κ, I*_*ν*_(*κ*) overflows quite rapidly with *κ*. To avoid numeric overflow, we use the exponentially scaled modified Bessel function *e*^−*κ*^*I*_*ν*_(*κ*) in calculations (the scaling is motivated by the asymptotic expansion of *I*_*ν*_(*κ*) ∼ *e*^*κ*^(2*πκ*)^−1/2^Σ_*t*_α_*t*_(*ν*)*κ*^−*t*^ for *κ* → ∞ (Abramowitz and Stegun, 1965)). The first order derivative of the exponentially scaled modified Bessel function is

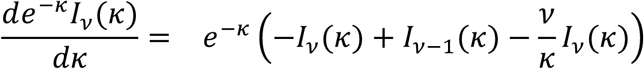

Previous work has used vMF distribution as the latent distribution for variational auto-encoders (Davidson et al., 2018; Guu et al., 2018; Xu and Durrett, 2018), but only the spherical variational auto-encoder (Davidson et al., 2018) learns the concentration parameter *κ*.

### 4.4 Sampling from a vMF distribution

Samples from vMF distributions can be obtained through a rejection sampling scheme (Ulrich, 1984; Wood, 1994). The algorithm is based on the theorem (Ulrich, 1984) that a *M*-dimensional vector 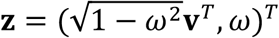 has a vMF distribution with direction vector (0, …, 1)^*T*^ ∈ 𝕊^*M*−1^ and concentration parameter *κ* if *ω* has a univariate density function with the following density function:

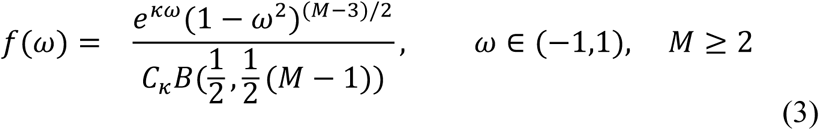

where **v** is uniformly distributed in 𝕊^*M*−2^, *C*_*κ*_ is a normalization term such that *f*(*ω*) is a legitimate density function, and 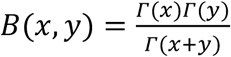 is the Beta function. The vector **v** is uniformly distributed in 𝕊^*M*−2^ and can be sampled from a standard normal distribution in *M* − 1 dimensions and then we normalize the resulting sample to unit length.

We then use rejection sampling to sample *ω* from the univariate distribution in **Equation 3**. The envelope function used for rejection sampling is defined as

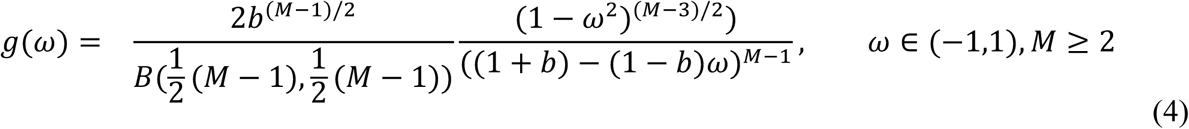

Where the term (Hornik and Grün, 2014) 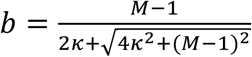. To sample from *g*(*ω*), we can first sample 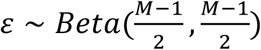 and pass the sample *ε* to the invertible function 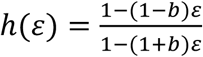. We can easily prove that 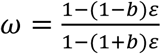 is distributed according to **Equation 4** based on the rule of transforming a continuous random variable with an invertible function. A sample *ω* is accepted if *κω* + (*M* − 1)log(1 − *x*_0_*ω*) − *c* ≥ log(*u*), where 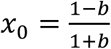, and 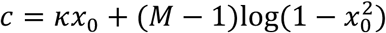 and *u* is sampled from a continuous uniform distribution with support in (0,1). The vector 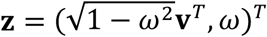 is a sample from vMF(**z**′ | **e**_1_, *κ*), where **e**_1_ = (0, …, 1)^*T*^ ∈ 𝕊^*M*−1^. We can then rotate **z**′ using a Householder matrix **I** − ***uu***^**T**^ to get a sample from vMF((**z** | **μ**, *κ*) (Davidson et al., 2018), where **I** is the identify matrix of rank *M* and 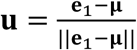, where ‖·‖ is the Euclidean norm. Overall, the samples from a Beta distribution are transformed and accepted or rejected by the rejection sampling scheme, and combined with samples **v** from a uniform distribution in 𝕊^*M*−2^. The combined samples are further transformed to generate samples from the desired vMF distribution. Remarkably, previous work has shown that this reparameterization approach still holds for these samples (Davidson et al., 2018), and can be used to optimize the vMF parameters **μ** and *κ*, which are the outputs of the inference neural network (encoder).

For visualization purposes, we typically set *M* = 3. Then the univariate density function becomes 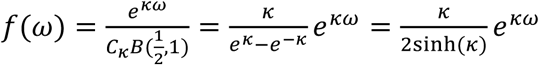, where sinh(·) is the hyperbolic sine function. We can directly draw samples from this density function by transforming a sample *ξ*, generated from a continuous uniform distribution *ξ* ∼ Unif(0,1) using the inverse cumulative function 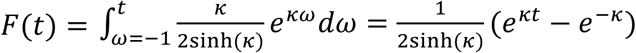. Specifically, we can use the following algorithm to generate a sample from *f*(*ω*):

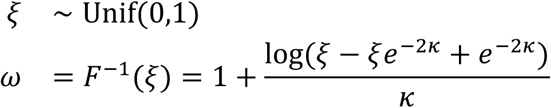

### 4.5 Poincaré ball and Lorentz model of the hyperbolic space

The Poincaré ball model represents the hyperbolic space as the interior of a unit ball in the Euclidean space: ℙ = {**z** ∈ ℝ^*d*+1^ |‖ **z** ‖< 1, *z*_0_ = 0, *d* ∈ ℤ^+^}, where **z** = (*z*_0_, …, *z*_*d*_)^*T*^. The distance between two points **z**_1_, **z**_2_ ∈ ℙ is defined as:

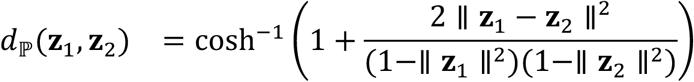

where 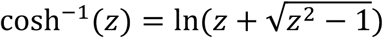 is the inverse hyperbolic cosine function, which is monotonically increasing for *z* ≥ 1. The symbol ‖·‖ represents the Euclidean norm. Notice that 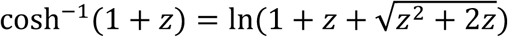, which approximates 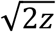 when lim *z* → 0 and ln(2*z*) for lim *z* → +∞. When both **z**_1_ and **z**_2_ are close to the origin with zero norm, *d*(**z**_1_, **z**_2_) ≈ cosh^−1^(1 + 2 ‖ **z**_1_ − **z**_2_ ‖^2^) ≈ 2 ‖ **z**_1_ − **z**_2_ ‖. Therefore, the Poincaré ball model resembles Euclidean geometry near the center of the unit hyperball. The induced norm of a point **z** ∈ ℙ is

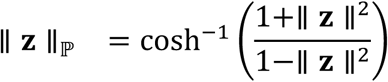

As **z** moves aways from the origin and approaches the border with ‖ **z** ‖≈ 1, the induced norm ‖ **z** ‖_ℙ_ grows exponentially. Hyperbolic geometry is useful to represent data with an underlying approximate hierarchical structure.

The Lorentz model is a model of the hyperbolic space and points of this model satisfy ℍ^d^ = {**z** ∈ ℝ^d+1^ | z_0_ > 0, ⟨**z, z**⟩_ℍ_ = −1}, where 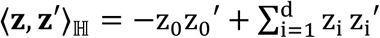 is the Lorentzian inner product (or Minkowski inner product when **z** ∈ ℝ^4^). The special one-hot vector **μ**_0_ = (1,0, …, 0)^T^ is the origin of the hyperbolic space. The distance between two points in the Lorentz model is defined as:

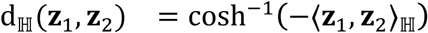

The tangent space of ℍ^d^ at point **μ** ∈ ℍ^d^ is defined as 𝒯_**μ**_ℍ^d^ ≔ {**z** | ⟨**μ, z**⟩_ℍ_ = 0}, *i.e.*, all the points that are orthogonal to **μ** based on the Lorentzian inner product. A point (z_0_, z_1_, …, z_d_)^**T**^ in the Lorentz model can be conveniently mapped to the Poincaré ball (Nickel and Kiela, 2018) for visualization:

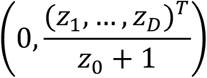

We discard the first element as it is a constant of zero.

### 4.6 Sampling from wrapped normal distributions of the Lorentz model

We used wrapped normal priors and wrapped normal posteriors defined in the Lorentz model to embed cells to a hyperbolic space (Grattarola et al., 2019; Mathieu et al., 2019; Nagano et al., 2019). A wrapped normal distribution in ℍ^*d*^ is constructed by first defining a normal distribution on the tangent space 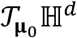 (an Euclidean subspace in ℝ^*d*+1^) at the origin **μ**_0_ = (1,0, …, 0)^*T*^ of the hyperbolic space. Samples from a normal distribution on the tangent space are transported to desired locations and further projected onto the final hyperbolic space (Nagano et al., 2019).

We used a set of invertible functions to transform samples from a normal distribution 𝒩(**z** | 0, **I**_*d*_**σ**) in ℝ^*d*^ to samples from a wrapped normal distribution in ℍ^*d*^ with mean of **μ**, where **σ** ∈ ℝ^*d*^ is the standard deviation of components *z*_1_ to *z*_*d*_, respectively, and **I**_*d*_ is the identity matrix in ℝ^*d*^ (Nagano et al., 2019; Rezende and Mohamed, 2015). First, let **z**_0_ = (0, **z**_0_′)^*T*^, which can be considered as a sample from 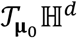, where **z**_0_′ is sampled from 𝒩(**z** | 0, **I**_*d*_**σ**). Next, **z**_0_ is parallel-transported to the tangent space 𝒯_**μ**_ℍ^*d*^ at **μ**, with coordinate **z**_1_, such that **z**_1_ is parallel to **z**_0_ (*i.e.*, pointing in the same direction relative to the geodesic) and preserves the norm (*i.e.*, ⟨**z**_0_, **z**_0_⟩_ℍ_ = ⟨**z**_1_, **z**_1_⟩_ℍ_) (Bergmann et al., 2018; Nagano et al., 2019):

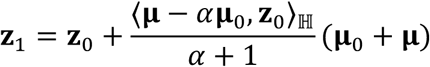

Finally, the exponential map (Grattarola et al., 2019; Nagano et al., 2019; Nickel and Kiela, 2018) projects **z**_1_ in the tangent space 𝒯_**μ**_ℍ^*d*^ back to the hyperbolic space by:

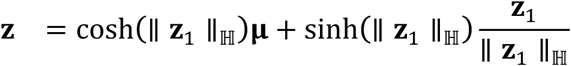

without altering the distance, *i.e., d*_ℍ_ (**μ, z**) = *d*_ℍ_ (**z**_1_, **z**_1_).

The likelihood after the invertible transformations can be calculated by 

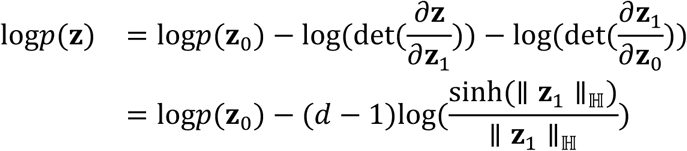

### 4.7 Model structure

As single cell data are sparse, with typically more than 90% genes with zero counts in each cell, we used softmax as the activation function to estimate the means of the negative binomial distributions and help generate sparse outputs from the decoders. We used the exponential linear units (ELU) (Clevert et al., 2015) activation functions for hidden layers, as it has been shown to improve convergence of stochastic gradient optimizations.

For all experiments, we used a three-layered encoder network (128-64-32) and a two-layered decoder network (32-128). We used the Adam stochastic optimization (Kingma and Ba, 2014) algorithm with a learning rate of 0.001. Only when embedding cells in more than 2D hyperbolic spaces, we used a learning rate of 0.0001 for numeric stability. For datasets with less than 10,000 cells, we trained models for 2,000 epochs. For datasets with more than 10,000 cells but less than 100,000 cells, we trained models for 500 epochs, and for the large number of immune cells with more than 2000,000 cells, we trained models for 250 epochs.

### 4.8 Visualization

The embeddings were visualized on a 3D sphere using the rgl package (Adler et al., 2003) from R, with the interactive 3D scatter plots saved as web graphics library files that can be opened in a browser. The rgl package uses OpenGL as the rendering backend, and can be used to rapidly and interactively visualize 3D scatter plots with millions of cells in a browser.

### 4.9 Data and code availability

We used publicly available datasets in this study. To make the results presented in this study reproducible, all processed data are available in the Single Cell Portal https://singlecell.broadinstitute.org/single_cell/study/SCP551/scphere#study-download The scPhere software package, implemented in TensorFlow, is available freely from https://github.com/klarman-cell-observatory/scPhere

### 4.10 Data

#### Cord blood mononuclear cells

This dataset (Stoeckius et al., 2017) consists of 8,617 cells, including 8,009 cord blood mononuclear cells and 608 mouse 3T3 fibroblasts, produced by the CITE-seq protocol (Stoeckius et al., 2017) on the 10x Chromium (v2) platform (Zheng et al., 2017). We only used the 2,293 CD14^+^ monocytes and the first 10 erythrocytes in the dataset. Based on the Seurat (Butler et al., 2018) tutorial (https://satijalab.org/seurat/v3.0/multimodal_vignette.html), we used the 2,000 highly variable genes in this study.

#### Cells of the colon mucosa

The cells were from the colon mucosa of 68 biopsies, collected from 18 ulcerative colitis patients and 12 healthy individuals (Smillie et al., 2019), and profiled by 10x Chromium (either v1 or v2). After filtering likely low-quality cells (clusters), we obtained a total of 301,749 cells (26,678 stromal cells and glia, 64,457 epithelial cells, and 210,614 immune cells as annotated in the original study (Smillie et al., 2019)). The cells span 12 stromal cell types/states, 12 epithelial cell types/states, and 23 immune cell types/states, identified by unsupervised clustering and manual annotations (Smillie et al., 2019). We used Seurat to select 1,307, 1,361, and 1,068 highly variable genes for the three major cell types, respectively, for scPhere analyses.

#### Human splenic NK cells

Cells were from a study profiling human and mouse splenic and blood NK cells (Crinier et al., 2018), and profiled by 10x Chromium (v2). We used the 1,755 human splenic NK cells from donor one in this study. We selected 2,724 highly variable genes and partitioned the 1,755 cells into four groups, labeled them as hNK_Sp1, hNK_Sp2, hNK_Sp3, hNK_Sp4, as in the original study (Crinier et al., 2018).

#### Human lung cells

Cells were from human lung tissue from asthma patients and healthy controls (Braga et al., 2019), and profiled by either 10x Chromium or Drop-seq (Macosko et al., 2015). We used the 3,314 cells from a donor prepared by the Drop-seq protocol that can be accessed from GEO: GSE130148.

#### Mouse white adipose tissue stromal cells

This dataset contains 1,378 cells from mouse white adipose tissue (Hepler et al., 2018) profiled by 10x Chromium (v2). In the original study, the authors only analyzed 1,045 tdTomato-mGFP+ cells and identified adipocyte precursor cells (APC), fibro-inflammatory progenitors (FIP), committed preadipocytes, and mesothelial cells. We analyzed all the cells and further identified pericytes, macrophages, and two groups of doublets.

#### *C. elegans* embryonic cells

This dataset consists of 86,024 *C. elegans* embryonic cells (Packer et al., 2019) profiled using the 10x Chromium (v2). The embryo times were partitioned into 12-time bins, and 63.5% of the cells were assigned to 36 major cell types based on annotation from GEO: GSE126954.

## Acknowledgements

We thank Jennifer Rood for helpful comments and Leslie Gaffney for help with figure preparation. Work was supported by the Klarman Cell Observatory, HHMI, the Food Allergy Science Initiative, Manton Foundation, the NIH BRAIN Initiative (1U19 MH114821), and NIH/National Institute of Diabetes and Digestive and Kidney Diseases grant (1RC2DK114784).

## Author contributions

J.D. and A.R. developed the model. J.D. conducted experimental analyses with guidance from A.R. J.D. and A.R. interpreted the results and wrote the manuscript.

## Competing interests

A.R. is a SAB member of ThermoFisher Scientific, Neogene Therapeutics, Asimov and Syros Pharmaceuticals. A.R. is a cofounder of and equity holder in Celsius Therapeutics and an equity holder in Immunitas.

## Supplementary Figure legends

**Supplementary Figure 1.**
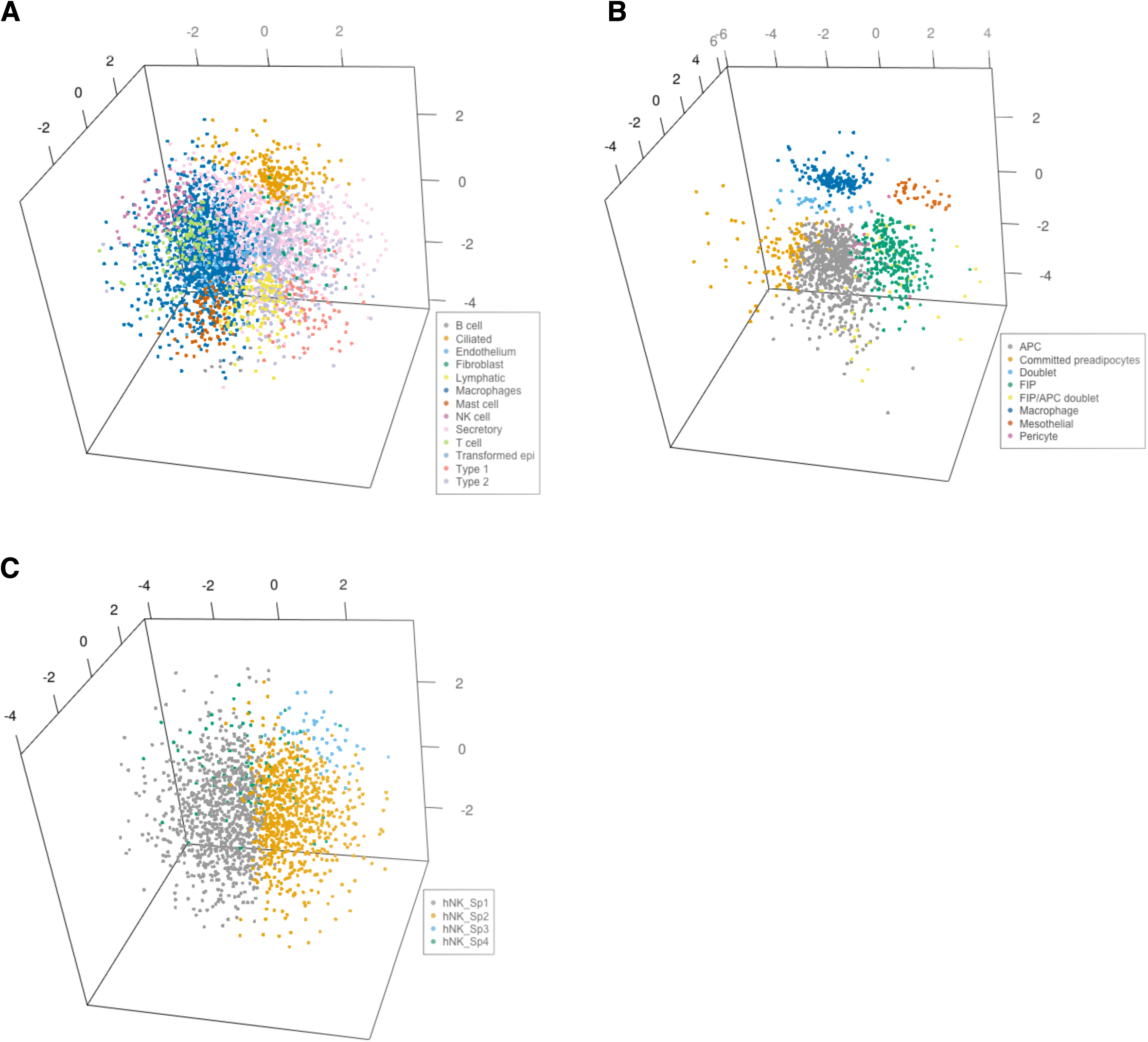
Embedding cells from different tissues in the 3D Euclidean latent space does not solve the cell crowding and occlusion problem. Embedding in the 3D Euclidean space of 3,314 human lung cells (**A**), 1,378 mouse white adipose tissue stromal cells (**B**), and 1,755 human splenic natural killer cells (**C**).

**Supplementary Figure 2.**
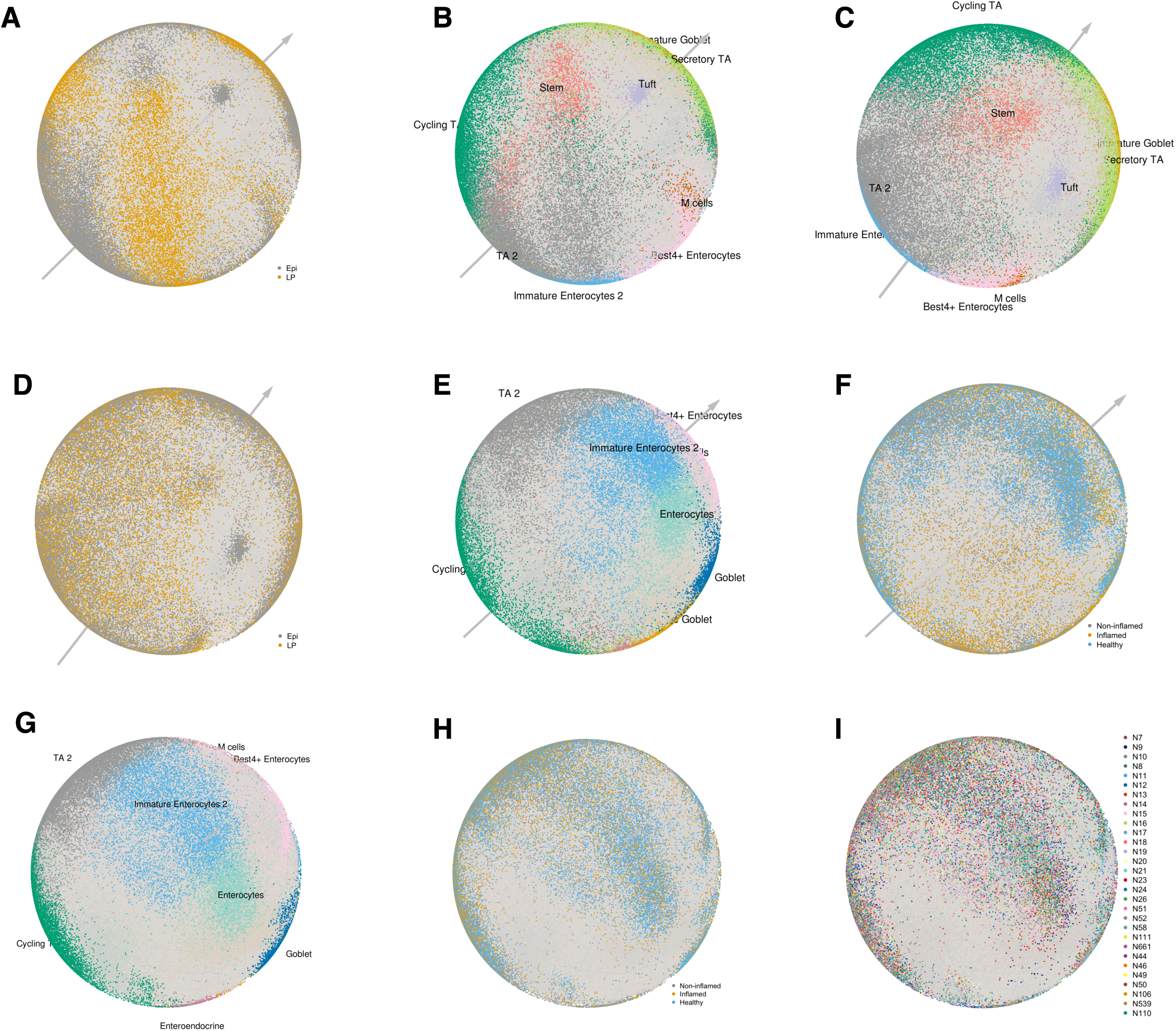
scPhere corrects multiple batch effects flexibly and efficiently. (**A-I**) Epithelial cell profiles from healthy individuals and UC patients embedded on the surface of unit spheres. (**A,B**) Taking only patient and disease status of each cell as the batch vector. Cells are colored by anatomical region (**A**) or type (**B**). (**C,D**) Taking the patient, disease status, and anatomical region as the batch vector. Cells are colored by type (**C**) or anatomical region (**D**). (**E,F**) Taking patient and anatomical region as the batch vector. Cells are colored by type (**E**) or disease status (**F**). (**G-I**) Taking the patient, disease status, and the anatomical region as the batch vector. This is a different view of the sphere in (**C,D**). Cells (dots) are colored by type (**G**), disease status (**H**), and patient (**I**).

**Supplementary Figure 3.**
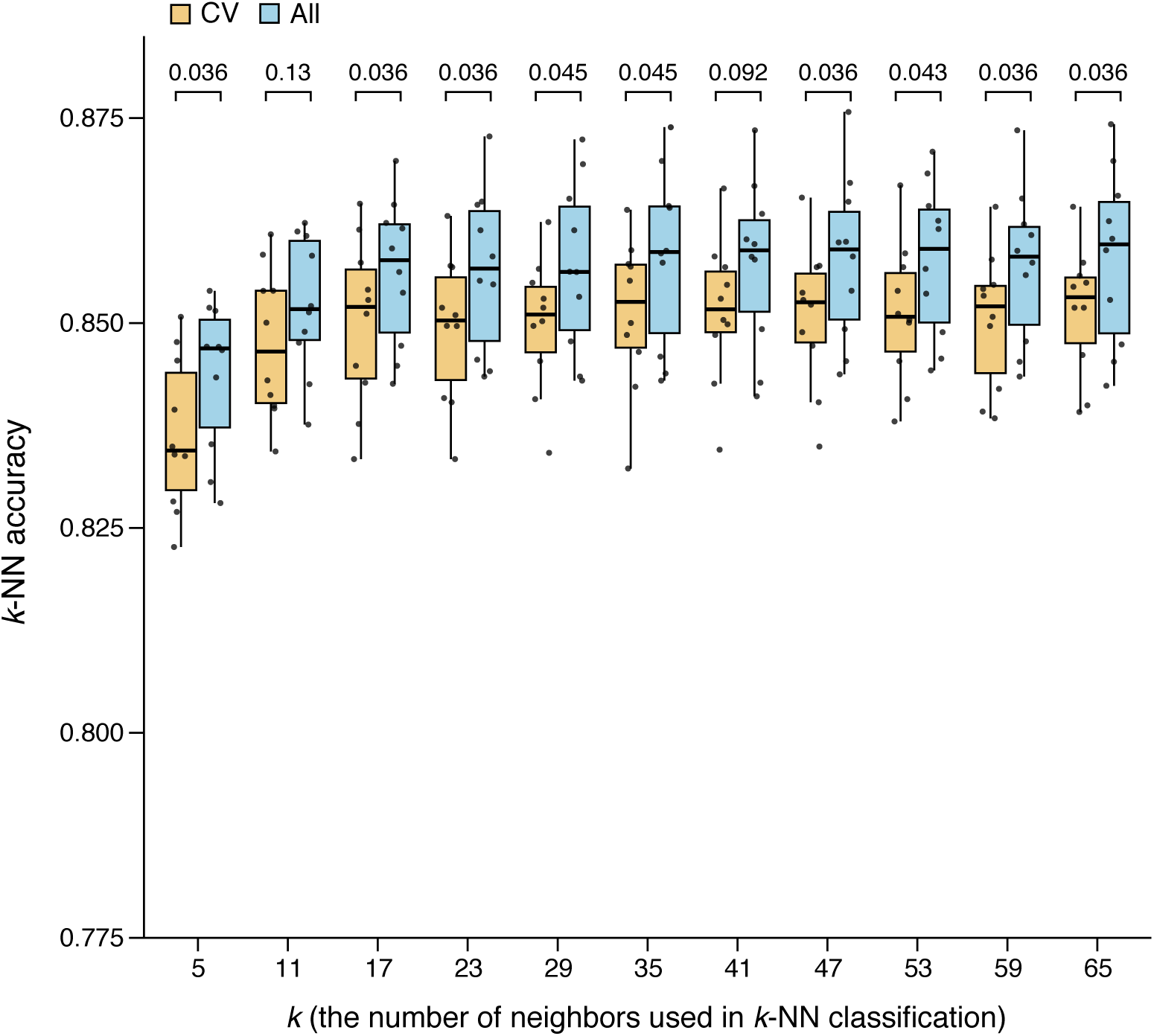
Training scPhere to classify cells from unseen test data. Classification accuracy (*y* axis) in 10-fold cross-validation *k*-nearest neighbor classification (*k*-NN) on 5D hyperspheres for different values of *k* (*x* axis) for test data classified either from classifiers trained on data mapped on a 5D hypersphere (orange; variable gene selection, scPhere modeling fitting, and J-NN classifier training were done in each fold) or when the 5D representation of cells was pre-computed by learning a scPhere model using all the cells (blue). Adjusted p-values (FDR, paired Wilcoxon rank sum test, two-sided) comparing the classification accuracies are at the top. Boxplots denote the medians and the interquartile ranges (IQRs). The whiskers of a boxplot are the lowest datum still within 1.5 IQR of the lower quartile and the highest datum still within 1.5 IQR of the upper quartile.

**Supplementary Figure 4.**
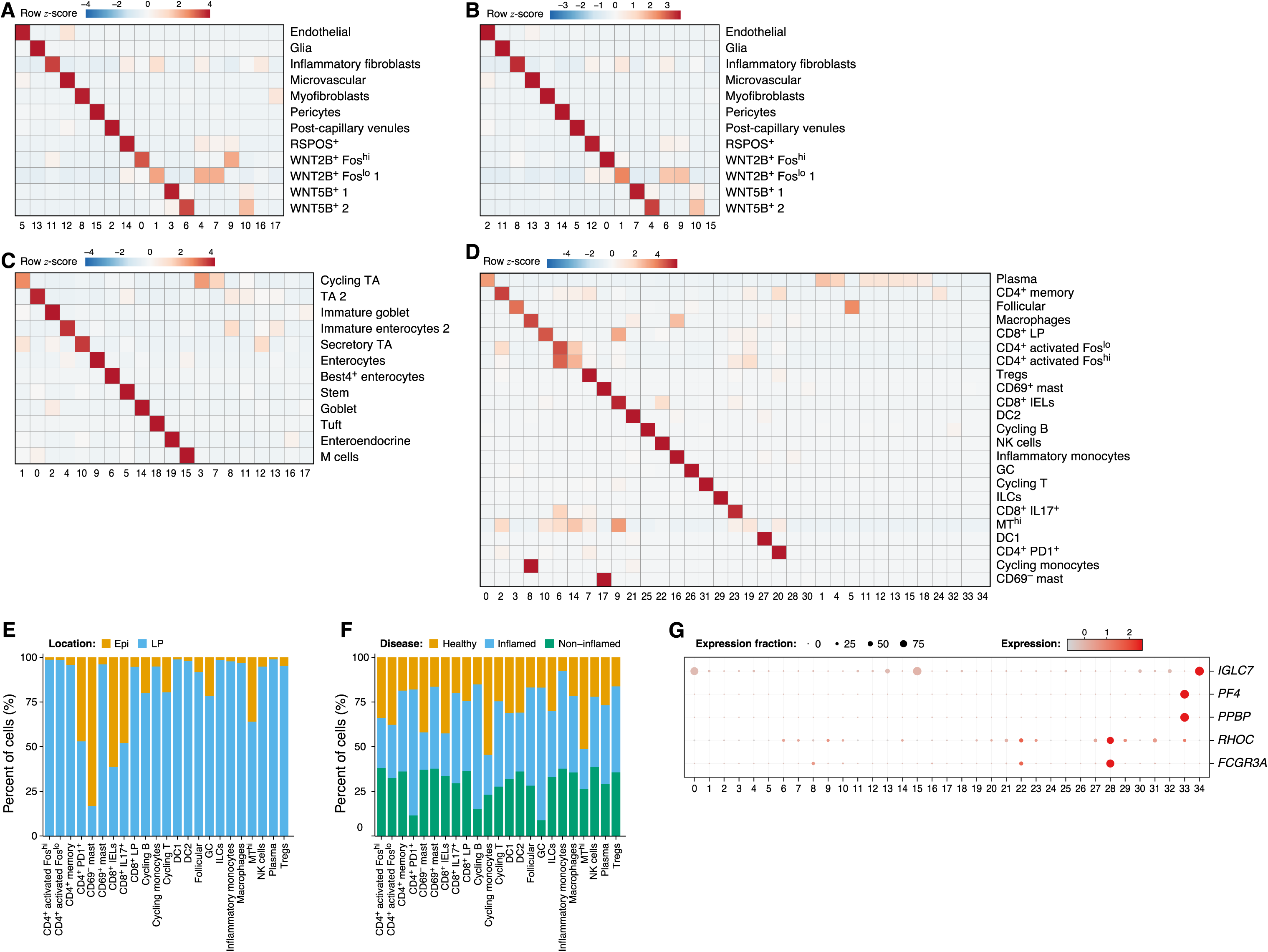
Clustering colon mucosa cells in the latent representations of 5D hyperspheres. (**A-D**) Comparison of cell assigned to Louvain clusters obtained on the cell embedding to a hypersphere (columns) and in the original study (rows) for stroma cells with 5D (**A**) or 10D (**B**) hypersphere embedding, epithelial cells (**C**) and immune cells (**D**). Color bar: z-scores (row-centered and scaled) of the number of cells. (**E,F**) Location and disease distribution vary across immune cell clusters. Percent of cells (*y* axis) from a given location (**E**, Epi or LP) or disease state (**F**, healthy, uninflamed, or inflamed) in each immune cell type annotated in the original study (*x* axis). (**G**) New immune clusters have distinctive expression markers. Fraction of expressing cells (dot size) and mean level of expression in expressing cells (dot color, row *z*-score of log(transcripts per 10,000 + 1)) for selected marker genes (rows) differentially expressed in small clusters of immune cells (cluster 28, 33, and 34, columns).

**Supplementary Figure. 5.**
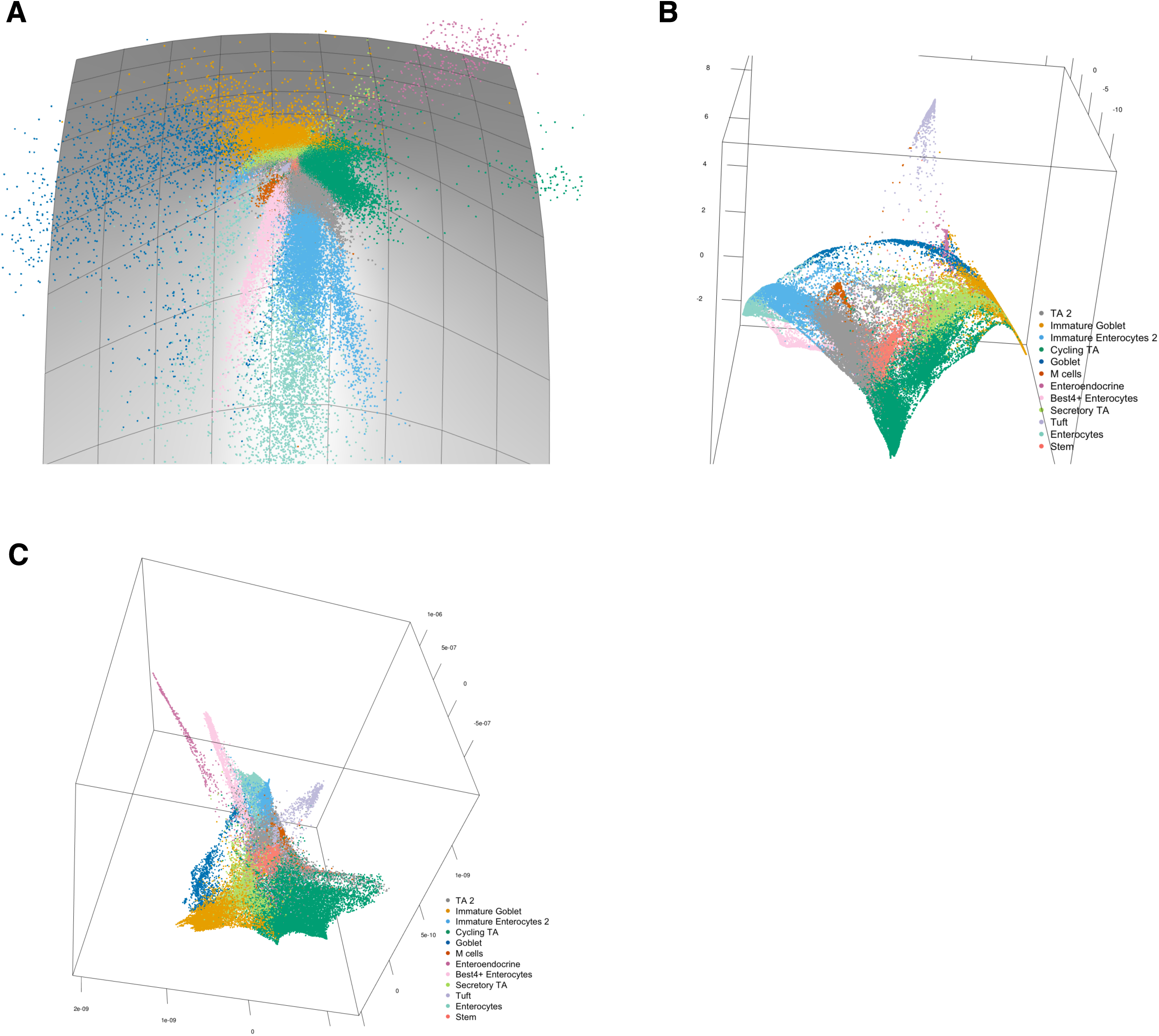
Embedding epithelial cells into a hyperbolic space. Embedding of epithelial cells colored by cell type in the 2D hyperbolic space of the Lorentz model (**A**), and PHATE multidimensional scaling with the 5D representations of cells in either the hyperbolic space of the Lorentz model (**B**) or in the Euclidean space (**C**) as inputs.

**Supplementary Figure 6.**
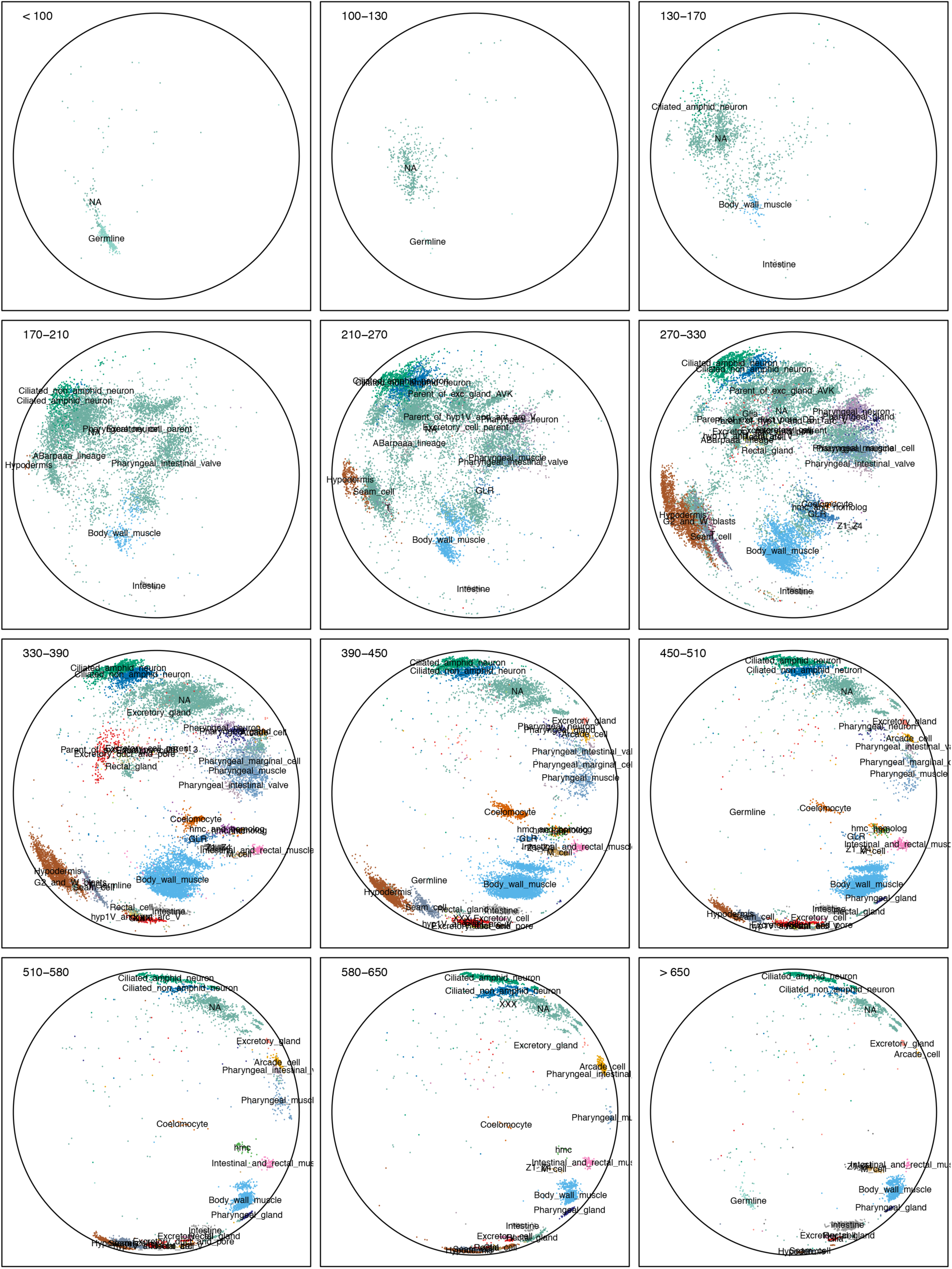
Poincaré disk embedding highlights the progression of *C. elegans* embryonic cells in time. Embedding of all *C. elegans* embryonic cells in a Poincaré disk (as in **Fig. 4C,E**), but each panel showing only the cells from one of 12 embryonic time bins (<100, 100-130, 130-170, 170-210, 210-270, 270-330, 330-390, 390-450, 450-510, 510-580, 580-650, >650). Cells are colored by annotated cell types (Packer et al., 2019).

**Supplementary Figure 7.**
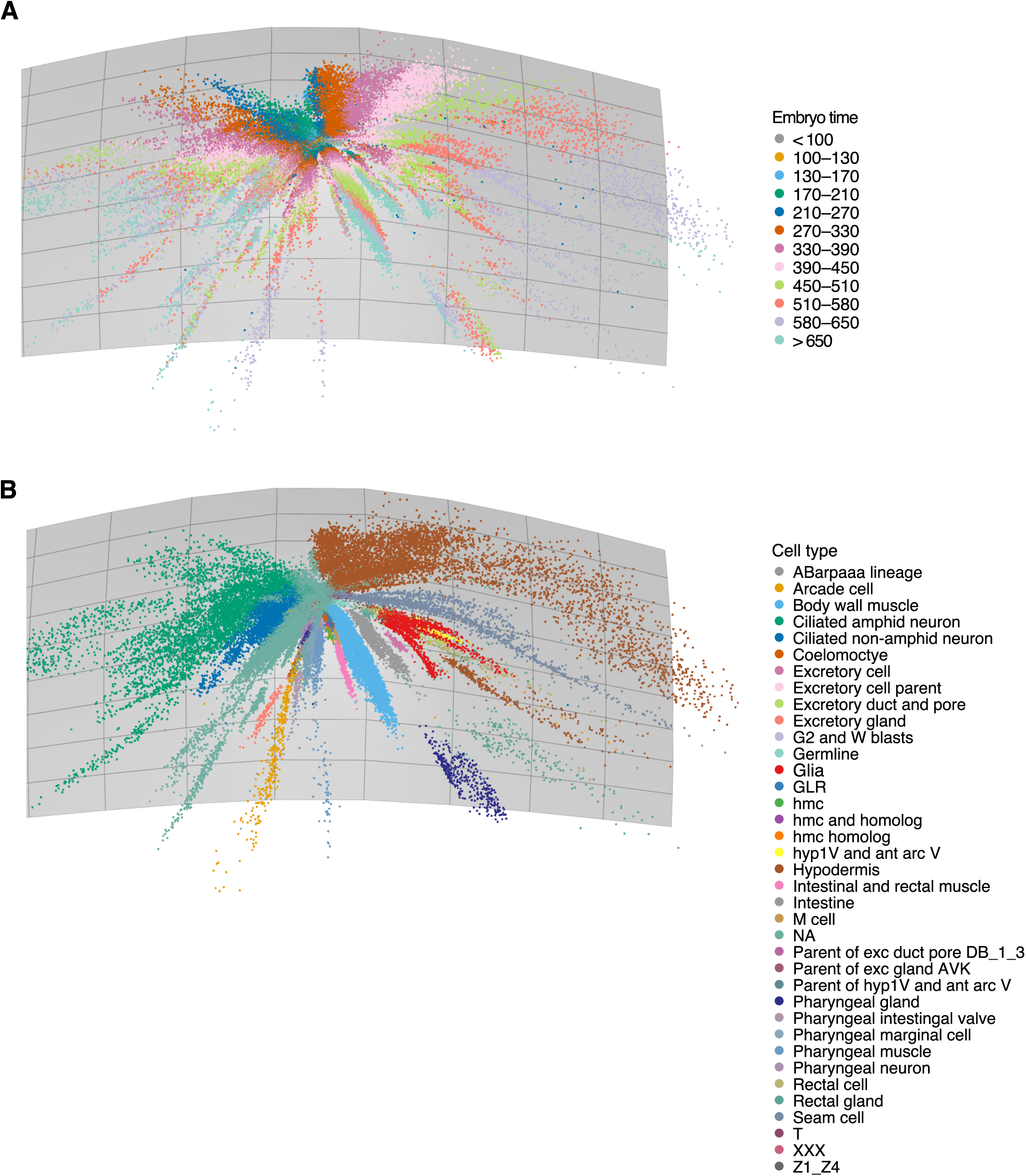
Embedding *C. elegans* embryonic cells into a hyperbolic space of the Lorentz model highlight differentiation branches. Embedding of *C. elegans* embryonic cells in the Lorentz model, colored by embryo time (**A**) or cell type (**B**).

**Supplementary Figure 8.**
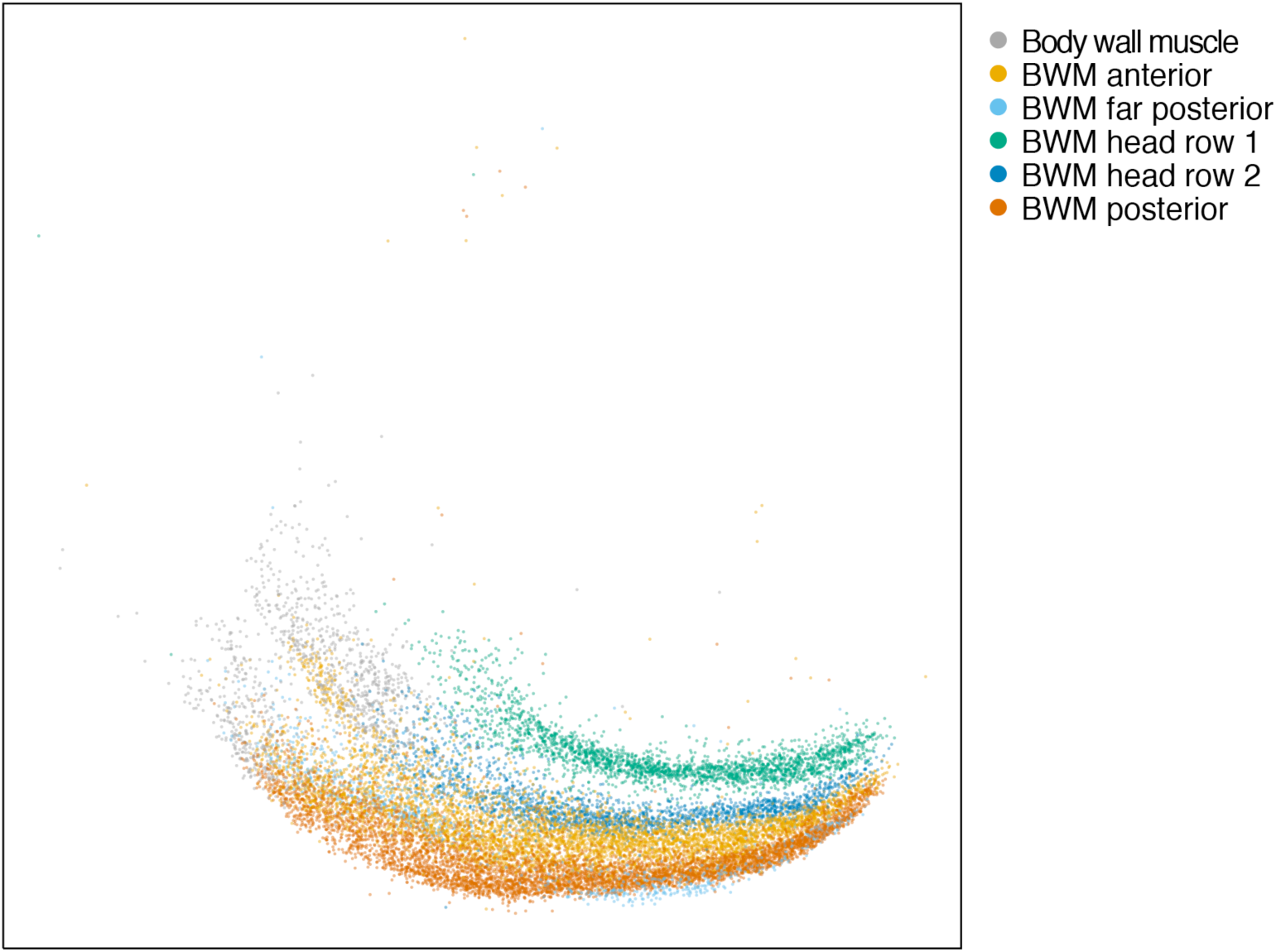
The *C. elegans* body wall muscle cells are ordered by lineages in the Poincaré disk. Embedding of *C. elegans* embryonic cells in a Poincaré disk (as in **Fig. 4C,E**), but showing only body wall muscle (BWM) cells, colored by lineage/location from first row and second row BWMs (D and MS lineage) to posterior BWMs (C lineage).

## Supplementary Movies

**Supplementary Movie 1. Embedding of stromal cells from human colon mucosa on the surface of the unit sphere, taking patient as the batch vector.**

Cells (dot) are color-coded by type. Annotations are marked adjacent to the corresponding cells.

**Supplementary Movie 2. Embedding stromal cells from human colon mucosa on the surface of the unit sphere, taking both patient and disease as the batch vector.**

Cells (dot) are color-coded by type. Annotations are marked adjacent to the corresponding cells. We did not include anatomical regions in the batch vector, because most cells were from the lamina propria fraction.

**Supplementary Movie 3. Embedding epithelial cells from human colon mucosa on the surface of the unit sphere, taking patient, disease status, and anatomical region as the batch vector.**

Cells (dot) are color-coded by type. Annotations are marked adjacent to the corresponding cells.

**Supplementary Movie 4. Embedding all cells from human colon mucosa on the surface of the unit sphere, taking patient, disease status, and anatomical region as the batch vector.**

Cells (dot) are color-coded by type. Annotations are marked adjacent to the corresponding cells.

**Supplementary Movie 5. Clustering the 5D latent representations of immune cells from human colon mucosa.**

Embedding immune cells on the surface of a unit sphere, taking patient, disease status, and anatomical region as the batch vector. Cells (dot) are color-coded by cluster membership numbers (and assigned cell types) as determined by Louvain clustering (resolution=1.2, the number of nearest neighbors=25) of the latent representations from embedding the cells on the surface of a 5D hypersphere.

## Notes

https://github.com/klarman-cell-observatory/scPhere

